# Transcriptional profiling of Hutchinson-Gilford Progeria patients identifies primary target pathways of progerin

**DOI:** 10.1101/2025.09.18.677125

**Authors:** Sandra Vidak, Sohyoung Kim, Tom Misteli

## Abstract

Hutchinson Gilford Progeria Syndrome (HGPS) is an ultra-rare pediatric premature aging disorder. The disease is caused by a point mutation in the *LMNA* gene leading to the production of the dominant-negative progerin isoform of the nuclear envelope protein lamin A. Disease severity and progression amongst the population of ∼140 known patients is variable. Most of the mechanistic insights into the disease have come from studies using cellular or mouse models of HGPS. To probe the clinical relevance of previously implicated cellular pathways and to address the extent of gene expression heterogeneity between patients, we have performed transcriptomic analysis of a comprehensive set of HGPS patients. We find misexpression of several cellular pathways across the patient population, particularly of multiple signaling pathways as well as the Unfolded Protein Response (UPR) and mesodermal cell fate specification. Variability amongst individual patients was limited, with misregulation of the major pathways observed in most patients. Comparing the transcriptome of patients with an inducible HGPS cell model, we distinguished immediate-early cellular response pathways from secondary adaptive pathways and identified mTORC1, the UPR, UV response, apoptosis and TNFα signaling via NF-κB as primary targets of the disease-causing progerin protein.

## INTRODUCTION

Hutchinson-Gilford Progeria Syndrome (HGPS) is an extremely rare genetic disorder characterized by segmental premature aging with an incidence of approximately 1 in 4–8 million live births [1, 2]. Although infants appear phenotypically normal at birth, clinical manifestations typically emerge between 9 and 12 months of age and include growth retardation, short stature, alopecia, joint contractures, osteolysis, and progressive lipodystrophy [1, 3–5]. As the disease progresses, patients experience arthritis and cardiovascular pathology, including accelerated atherosclerosis and arterial stiffening, with death typically occurring from myocardial infarction or stroke around a mean age of 14.5 years [6–9].

Classic HGPS is caused by a *de novo* heterozygous point mutation (c.1824C>T; G608G) in exon 11 of the *LMNA* gene, which encodes the nuclear lamina proteins lamin A and C [10, 11]. Although the disease-causing mutation does not alter the predicted amino acid sequence of lamin A, it activates a cryptic splice donor site in the LMNA pre-mRNA, resulting in the production of a truncated lamin A isoform referred as progerin [10, 11]. This aberrant isoform lacks 50 amino acids at the C-terminus, including the cleavage site for the ZMPSTE24 metalloprotease, thereby retaining a farnesylated C-terminal motif that permanently anchors progerin to the inner nuclear membrane [2, 12, 13]. Progerin exerts dominant-negative effects that disrupt nuclear architecture and impair critical cellular functions. Its accumulation leads to nuclear blebbing, loss of heterochromatin, impaired DNA repair, altered gene expression, and defects in mechanotransduction [14–16]. Additionally, progerin aggregates at the nuclear periphery sequester essential regulatory proteins, such as the oxidative stress response factor NRF2 and molecular chaperones in the endoplasmic reticulum, further exacerbating cellular dysfunction [17–19].

Studies using cellular and mouse models of HGPS, including transcriptomic analysis [20–23], have implicated several major pathways as contributors to the disease phenotype [2, 13]. The most prominent of these pathways include altered signaling by nuclear factor kappa B (NF-kB) [24], transforming growth factor beta (TGF-β) [25], mammalian target of rapamycin (mTORC1) [26, 27], Wnt/b-catenin [28] and the unfolded protein response (UPR) [18, 29], alongside mitochondrial impairment [30–32]. HGPS cells also exhibit significant metabolic dysregulation, particularly in glucose and lipid metabolism [33], and impaired antioxidant responses due to NRF2 pathway dysfunction, leading to increased oxidative stress [17]. Importantly, stem cell exhaustion and compromised regenerative capacity further exacerbate tissue degeneration [34–36], partly by activation of the Notch signaling pathway [37]. Fibrotic remodeling and chronic inflammation are observed across multiple organ systems [38], contributing to functional decline, while vascular pathology, marked by atherosclerosis and valve calcification drives the cardiovascular complications that ultimately lead to early mortality [6]. Collectively, these interconnected pathways orchestrate the systemic premature aging phenotype observed in disease models. While some of these pathways have been found to be dysregulated in select patient-derived cell lines, it is unclear how prevalent the misregulation of these, and other, pathways is in the patient population. The degree of conservation of misregulated cellular pathways in the patient population is also of interest considering that HGPS patients often show highly variable disease symptoms and progression [8, 9, 39].

In this study, we conduct a comprehensive analysis of the transcriptional status of patient derived primary HGPS dermal fibroblasts and age-matched and/or parental control cell lines using RNA-seq analysis. We performed transcriptome analysis on fibroblast cell lines from 20 HGPS patients, representing about 15% of the world’s patient population. Transcriptomics identified several prevalent major misregulated pathways in HGPS patients, many of which are also affected in mouse or cellular models of the disease. Pathway analysis shows homogeneous deregulation of various pathways across the patient population. Comparison of patient transcriptomics with the effect of acute induction of progerin in a HGPS cell-based model allowed us to distinguish immediate early progerin-induced pathways from long-term adaptive pathways in patient cells. Pathway analysis identified mTORC1 signaling, the UV response, apoptosis, the UPR and TNFα signaling via NF-κB pathways as immediate targets of progerin, and oxidative phosphorylation, Notch signaling and Wnt/b-catenin signaling as adaptive or compensatory pathways affected by long term progerin expression in HGPS patients. These results explore the role of transcriptional heterogeneity in the HGPS patient population and identify clinically relevant primary target pathways of the disease-causing progerin protein.

## MATERIALS AND METHODS

### Cell culture

Primary human dermal fibroblast cell lines were obtained from the Progeria Research Foundation (PRF; https://www.progeriaresearch.org/cell-and-tissue-bank/) or the National Institute of Aging (NIA) Cell Repository distributed by the Coriell Institute. Ten cell lines were from healthy donors, sixteen were from patients that have the classic mutation in *LMNA* Exon 11, heterozygous c.1824C > T (p.Gly608Gly), and four were from patients with non-classic mutations. The list of the cell lines used together with the source and the age at donation are listed in Table 1. Cells were grown in high glucose DMEM containing 15% fetal bovine serum (FBS), 1% GlutaMAX (ThermoFisher #35050-061) and 1% Penicillin-Streptomycin (ThermoFisher #15140-122) at 37 °C in 5% CO_2_. All cell lines were mycoplasma negative as shown by routine testing (EZ-PCR™ Mycoplasma Detection Kit, Biological Industries). hTERT-immortalized GFP-progerin doxycycline inducible dermal fibroblast cell lines were maintained and induced for 6 days as described [40].

**Table 1.**
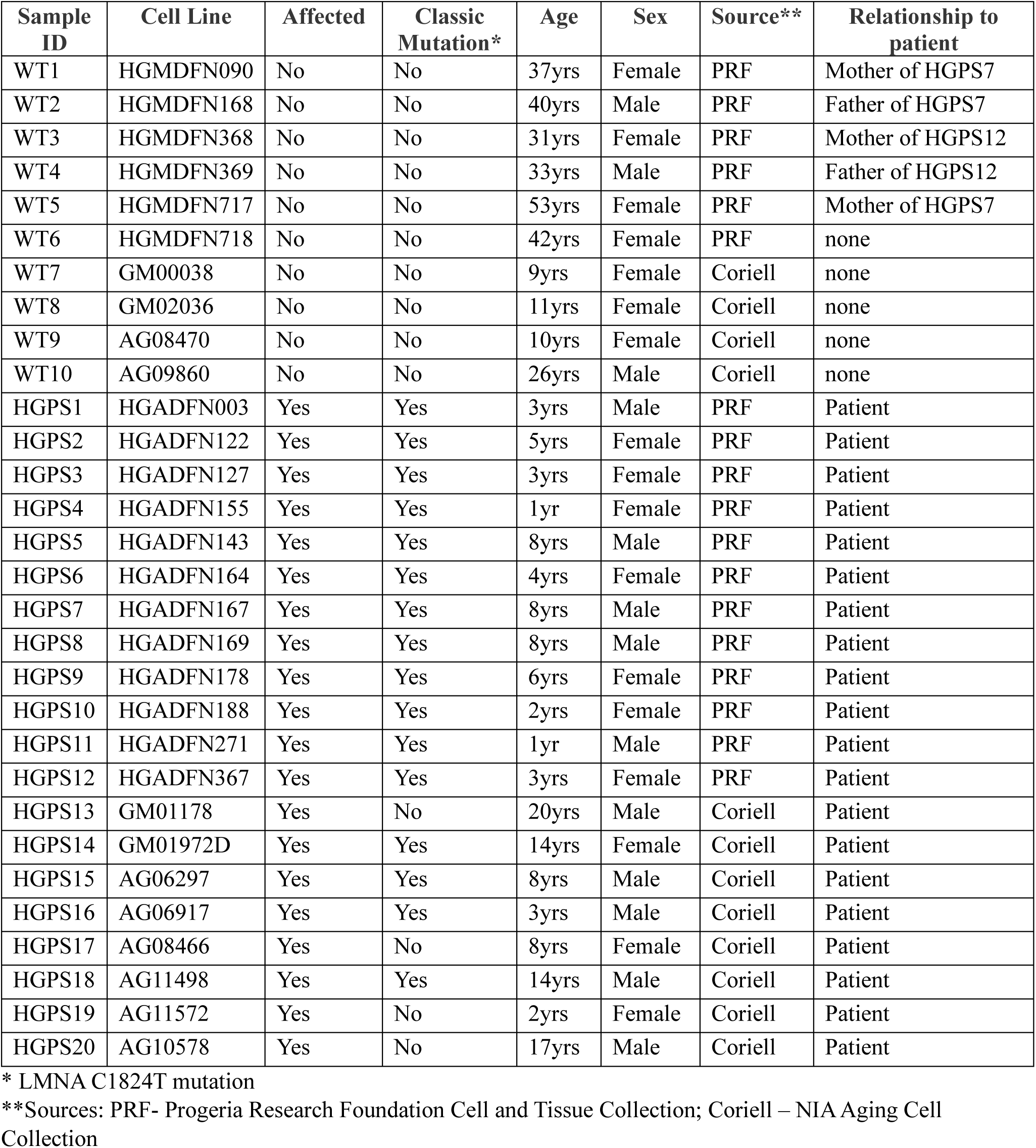
HGPS patients and healthy controls used in this study.

| **Sample ID** | **Cell Line** | **Affected** | **Classic Mutation\*** | **Age** | **Sex** | **Source\*\*** | **Relationship to patient** |
| --- | --- | --- | --- | --- | --- | --- | --- |
| WT1 | HGMDFN090 | No | No | 37yrs | Female | PRF | Mother of HGPS7 |
| WT2 | HGMDFN168 | No | No | 40yrs | Male | PRF | Father of HGPS7 |
| WT3 | HGMDFN368 | No | No | 31yrs | Female | PRF | Mother of HGPS12 |
| WT4 | HGMDFN369 | No | No | 33yrs | Male | PRF | Father of HGPS12 |
| WT5 | HGMDFN717 | No | No | 53yrs | Female | PRF | Mother of HGPS7 |
| WT6 | HGMDFN718 | No | No | 42yrs | Female | PRF | none |
| WT7 | GM00038 | No | No | 9yrs | Female | Coriell | none |
| WT8 | GM02036 | No | No | 11yrs | Female | Coriell | none |
| WT9 | AG08470 | No | No | 10yrs | Female | Coriell | none |
| WT10 | AG09860 | No | No | 26yrs | Male | Coriell | none |
| HGPS1 | HGADFN003 | Yes | Yes | 3yrs | Male | PRF | Patient |
| HGPS2 | HGADFN122 | Yes | Yes | 5yrs | Female | PRF | Patient |
| HGPS3 | HGADFN127 | Yes | Yes | 3yrs | Female | PRF | Patient |
| HGPS4 | HGADFN155 | Yes | Yes | 1yr | Female | PRF | Patient |
| HGPS5 | HGADFN143 | Yes | Yes | 8yrs | Male | PRF | Patient |
| HGPS6 | HGADFN164 | Yes | Yes | 4yrs | Female | PRF | Patient |
| HGPS7 | HGADFN167 | Yes | Yes | 8yrs | Male | PRF | Patient |
| HGPS8 | HGADFN169 | Yes | Yes | 8yrs | Male | PRF | Patient |
| HGPS9 | HGADFN178 | Yes | Yes | 6yrs | Female | PRF | Patient |
| HGPS10 | HGADFN188 | Yes | Yes | 2yrs | Female | PRF | Patient |
| HGPS11 | HGADFN271 | Yes | Yes | 1yr | Male | PRF | Patient |
| HGPS12 | HGADFN367 | Yes | Yes | 3yrs | Female | PRF | Patient |
| HGPS13 | GM01178 | Yes | No | 20yrs | Male | Coriell | Patient |
| HGPS14 | GM01972D | Yes | Yes | 14yrs | Female | Coriell | Patient |
| HGPS15 | AG06297 | Yes | Yes | 8yrs | Male | Coriell | Patient |
| HGPS16 | AG06917 | Yes | Yes | 3yrs | Male | Coriell | Patient |
| HGPS17 | AG08466 | Yes | No | 8yrs | Female | Coriell | Patient |
| HGPS18 | AG11498 | Yes | Yes | 14yrs | Male | Coriell | Patient |
| HGPS19 | AG11572 | Yes | No | 2yrs | Female | Coriell | Patient |
| HGPS20 | AG10578 | Yes | No | 17yrs | Male | Coriell | Patient |

\* LMNA C1824T mutation
\*\*Sources: PRF- Progeria Research Foundation Cell and Tissue Collection; Coriell – NIA Aging Cell Collection

### RNA isolation and RNA-seq sample preparation

Total RNA was extracted from cells using the NucleoSpin RNA Kit (Takara Bio) according to manufacturer instructions. The RNA 6000 nano assay on the Agilent bioanalyzer was used to quantitate the Total RNA samples. Stranded Total RNA Ligation with Ribo-zero plus library prep kit (Illumina) was used for sample preparation. Briefly, ribosomal RNA (rRNA) was removed using biotinylated, target-specific oligos combined with Ribo-Zero rRNA removal beads. The RNA was fragmented into small pieces and the cleaved RNA fragments copied into first strand cDNA using reverse transcriptase and random primers, followed by second strand cDNA synthesis using DNA Polymerase I and RNase H. The resulting double-strand cDNA was used as the input to a standard Illumina library prep with end-repair, adapter ligation and PCR amplification being performed to give a sequencing-ready library. The final purified product was quantified by qPCR before cluster generation and sequencing on a NovaSeq 6000 S4 flowcell. All samples were collected and analyzed in triplicates.

### RNA-seq preprocessing and analysis

Illumina stranded total RNA prepared libraries were pooled and sequenced on a NovaSeq 6000 S4 flowcell using 2×151 cycles paired-end sequencing. The Illumina RTA v3.4.4 was used to process raw data files and the Illumina bcl2fastq2.20 was used to demultiplex to generate sample specific fastq files. Samples had 140 to 270 million pass filter reads with more than 89% of bases above the quality score Q30. The sequencing reads were trimmed adapters and low-quality bases using Cutadapt (v 1.18). The trimmed reads were mapped to human reference genome (hg38) and the annotated transcripts (GENCODE v30) using STAR aligner (v2.7.0f) with two-pass alignment option.

### Transcriptome Analysis

Raw tag counts of exon regions at the gene level were obtained using the featureCount function in Subread (v 2.0.3) based on GENCODE annotation files. The raw tag counts were normalized using the default size factor in DESeq2 (v 1.42.1). Differential gene expression between two contrasting groups was evaluated using the Wald-test (FDR < 0.05) and |log2 fold change (LFC)| > log2(1.5) implemented in DESeq2. Four contrasting analyses were performed: individual HGPS vs. WT, group-level HGPS vs. WT, individual WT vs. all other WT, and GFP-progerin cell line against uninduced GFP-progerin cell line as illustrated in Figure 2B and Figure S3. For each HGPS patient, pairwise contrasts were performed against all 10 WT samples (e.g., HGPS1 vs. WT1, HGPS1 vs. WT2,…, HGPS1 vs. WT10), generating 10 log2 fold change estimates per HGPS patient. The mean of these 10 estimates was considered as the overall difference for that HGPS patient relative to WT (individual HGPS vs. WT). This procedure was repeated across all 16 HGPS samples, and the mean of these 16 values (HGPS1 vs. WT, HGPS2 vs. WT, HGPS3 vs. WT,…) represented the group-level HGPS vs. WT difference. Similarly, to estimate within WT variability, for each WT, the expression difference was calculated against all other WT samples (excluding itself). The individual and average gene expression differences were estimated within the framework of a generalized linear model in DESeq2. For pathway analysis, genes were ranked by Wald statistic obtained using DESeq2 and analyzed for Gene Set Enrichment Analysis (GSEA) using fgsea (v 1.28.0) with msigdb.v2023.2.Hs.symbols.gmt database downloaded from the GSEA website [PMID16199517]. Heatmaps of z-scaled, variance-stabilized gene expression values were generated using the pheatmap (v 1.0.12) with Eucliean distance and Ward.D2 clustering. All other heatmaps were generated using the default settings (Eucliean distance and complete linkage clustering method). All samples were analyzed in triplicates.

### Quantitation of splice site usage

The ratio of the progerin splice site usage was calculated as the proportion of reads supporting the progerin splice site relative to the total reads supporting either the progerin and wild-type Lamin A splice sites, using count_jct function in Alfred program (v 3.0.2) [PMID 30520945]. For progerin, sequencing tags supporting intra-gene exon-exon junctions were detected at chr1:156,138,607-chr:156,139,079 (99.6%) or chr1:156,138,607-chr:156,139,769 (21%) (hg38). For Lamin A, tags supporting intra-gene exon-exon junctions were detected at chr1:156138757-chr:156139079 (99.9%) or chr1:156,138,757-156,139,769 (0.1%) (hg38).

### Statistical Analysis

PCA of gene expression data was performed on variance-stabilized data obtained via vst function and plotPCA function in DESeq2 using the top 500 most variable genes across samples. PCA of Normalized Enrichment Score (NES) obtained from GSEA was performed using the svd function available in R (v 4.3.2) on mean-centered data.

### Data availability

The HGPS patient and healthy volunteer sequencing data reported in this paper were deposited on NCBI Gene Expression Omnibus (GEO) and are accessible through GEO accession number GSE306264.

## RESULTS

### Characterization of HGPS patient skin fibroblasts

To transcriptionally profile HGPS patients, we performed RNA sequencing (RNA-seq) analysis of a collection of primary patient-derived skin fibroblasts from twenty patients (ages 1-20 years) characterized as HGPS in publicly available cell banks from either The Progeria Research Foundation or the Coriell NIA Aging Cell Collection, and ten healthy control individuals from the same sources (ages 9-53 years) (Table 1) (see Materials and Methods). To confirm that all cell lines carry the classic 1824C>T HGPS mutation, we used the transcriptomics data to test for the presence of the mutation and the progerin transcript in all cell lines (Fig. 1 A-B; S1A; Table 1). Expression of the *LMNA* progerin transcript, containing the characteristic internal deletion of 150 nucleotides (Fig. 1A), was readily detected in sixteen of the twenty cell lines, but not in control samples (Fig.1C; S1B, full dataset available in GEO). Despite their HGPS classification in the Coriell repository, four patient samples (HGPS 13, 17, 19, 20) lacked the classic HGPS mutation in the *LMNA* gene and no progerin mRNA was detectable (Fig. 1 A-C; Fig.S1A-B; Table 1). The distinct nature of these patient cell lines was also evident when transcriptomic data were analyzed by principal component analysis (PCA) based on their general gene expression pattern. According to the first principal component, the four patient cell lines that lacked progerin transcripts segregated strongly from progerin-expressing HGPS samples (Fig. S1B). We conclude that these cell lines do not represent classic HGPS but rather are derived from patients with related progeroid syndromes and phenotypes. This classification is in line with the description of progeroid laminopathies which are caused by mutations in *LMNA* other than 1824C>T [7], as well as the occurrence of atypical progeroid syndromes caused by non-*LMNA* mutations [41, 42] (see Discussion). We consequently excluded the four samples lacking the classic *LMNA* 1824C>T mutation from further analysis. The remaining 16 progeria patient samples effectively segregated from healthy controls based on the first principal component (Fig.1D, Fig. S1C), with the exception of one HGPS patient (HGPS14) which clustered more closely with the control samples (Fig. 1D, S1C; see below).

**Figure 1.**
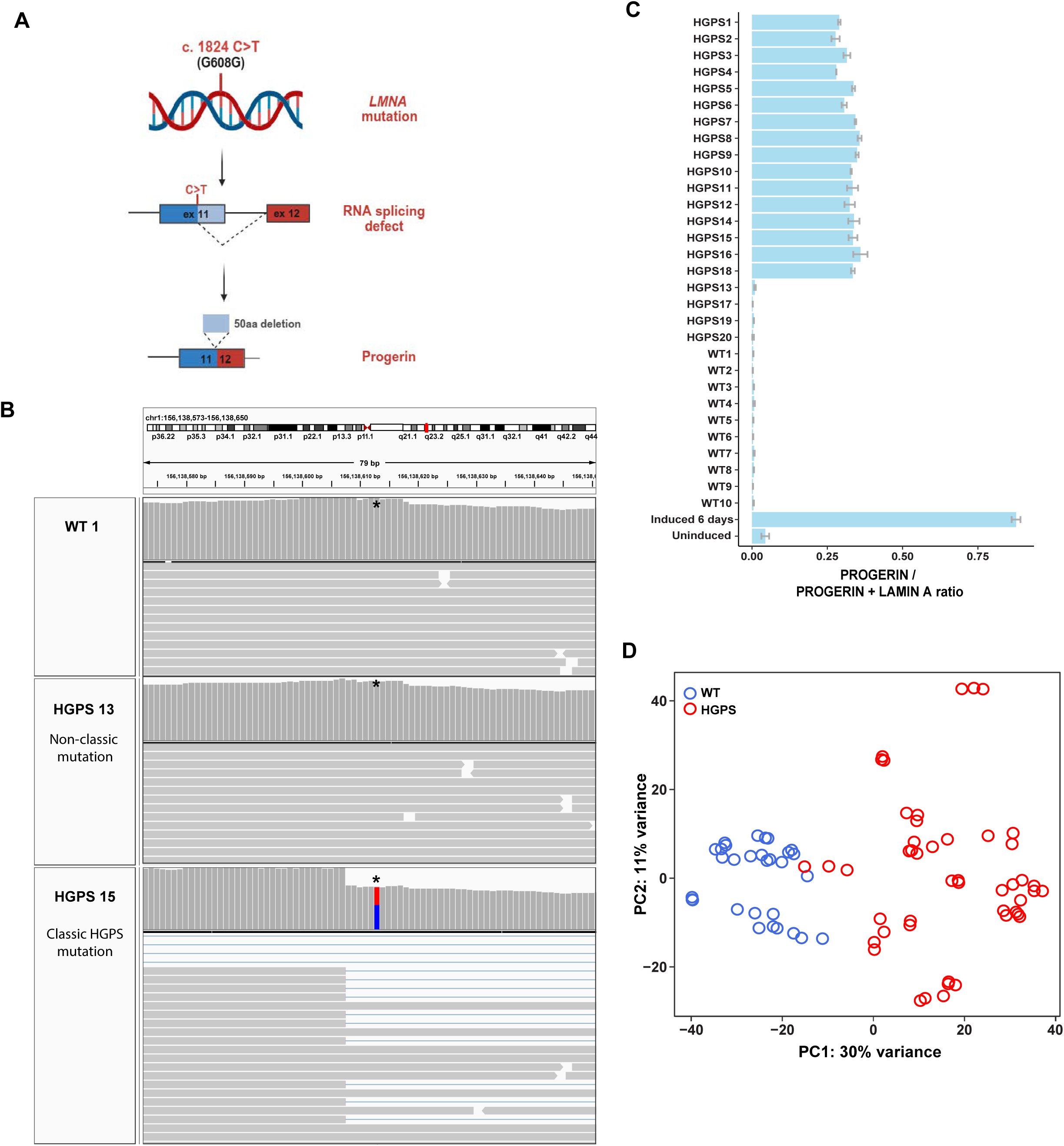
Characterization of primary human dermal fibroblasts. **(A)** A single point mutation c.1824C>T in exon 11 of the *LMNA* gene results in aberrant splicing and subsequent translation into a truncated lamin A protein isoform with an internal deletion of 50aa, termed progerin, which causes the classic form of progeria called Hutchinson Gilford Progeria Syndrome (HGPS). **(B)** Genome browser view showing the presence or absence of the classic c.1824C>T HGPS mutation in the *LMNA* gene of a healthy control and two progeria patients. One patient (HGPS15) carries the classic HGPS mutation (red/blue), the other (HGPS13) an atypical progeria lacking the c.1824C>T mutation. *denotes the location of residue 1824. Horizontal gray bars show reads aligned to the reference genome (chr1:156,138,573-156,138,650). Vertical gray bars show sequencing depth at each location. The red and blue bar indicates the presence of heterozygous variants at the indicated genomic position. **(C)** Quantification of progerin expression in transcripts from healthy controls, progeria patient samples and inducible GFP-progerin skin fibroblast cell line, shown as the ratio of progerin splice site usage over the usage of progerin and wild type lamin A splice sites. Error bars indicate the standard error of fraction values from three replicates. Note that HGPS samples 13, 17, 19, 20 do not express progerin and were excluded from RNA-seq analysis. **(D)** Principal component analysis (PCA) of gene expression profiles of the primary HGPS patient and control fibroblasts used in this study after the exclusion of the four non-typical progeria patients. Samples primarily segregate by progeria vs control phenotype. The top 500 genes with the highest variability across samples were used for the PCA plot. Each circle represents an individual cell line sample; all samples were analyzed in triplicate. Blue circles represent healthy wild type control samples and red circles represent HGPS patients.

### Deregulation of major cellular pathways in HGPS patients

To identify genes that exhibit significant changes in the expression levels between WT and HGPS skin fibroblasts, we performed differential gene expression (DGE) analysis. Group-level comparison between HGPS and WT samples revealed 3292 significantly differentially expressed genes, of which 1471 were upregulated and 1821 were downregulated in HGPS (Fig.2A; FDR cut off: 0.05, LFC cut off: log2(1.5)). To assess HGPS-dependent transcriptome changes in individual patients, we performed pairwise comparisons of each HGPS patient against all ten WT samples, generating ten gene expression fold-change estimates per gene. The mean of these estimates was considered as the overall difference for each HGPS patient relative to WT (Fig. 2B, left). This approach enabled the identification of differentially expressed genes (DEGs) for each HGPS patient relative to all the WT samples. We identified 693 DEGs consistently up- or down-regulated in thirteen or more patients (80%) (Fig. 2C). The number of DEGs varied from 2729-6208 in individual HGPS cell lines (Table 2). In addition, the average gene expression fold change in each individual WT cell line compared to all other WT samples was calculated to identify HGPS-independent transcriptome changes in individual WT samples (Fig. 2B, right). This analysis identified 693 DEGs which segregate HGPS patients from WT samples, with the exception of patient HGPS14 which displayed closer clustering with healthy volunteers (Fig. 2C), consistent with PCA analysis (Fig. 1D). WT samples from individuals over 20 years (20-53) segregated from young individuals but no age-based clustering was observed for controls or HGPS patients under the age of 20 (Fig. 2C).

**Figure 2.**
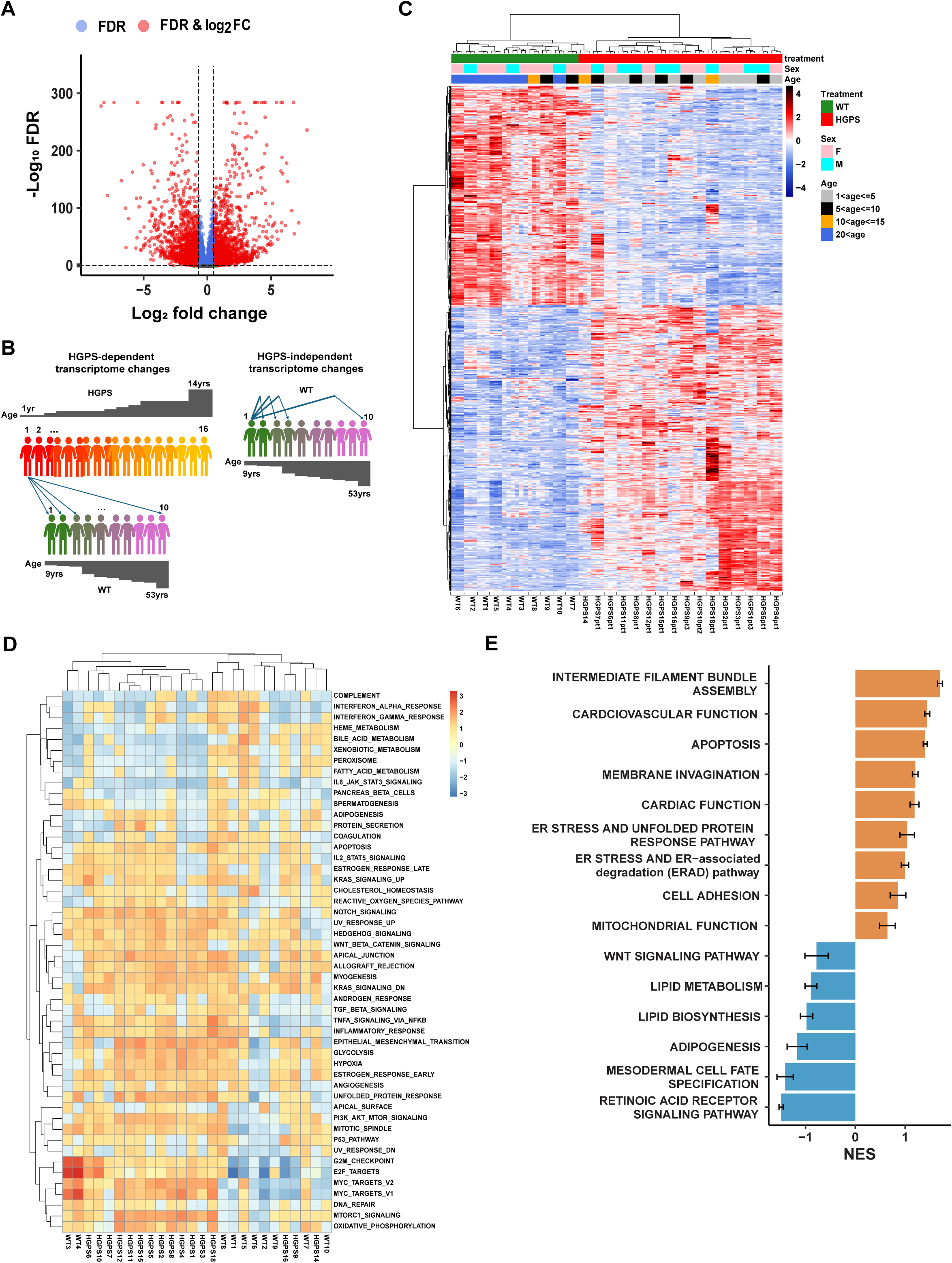
Misreguled cellular pathways in HGPS. **(A)** Volcano plot comparison of differentially expressed genes (DEGs) between WT controls and HGPS patients. FDR cut off: 0.05, LFC cut off: log2(1.5). Dashed lines indicate threshold (FDR < 0.05, |log2FC| > log2(1.5)). Data point colors denote whether the criteria for FC, FDR, or both were met. **(B)** Graphical representation of the DEG analysis in individual patients and healthy controls. For HGPS-dependent transcriptome changes we estimated average gene expression fold change in each HGPS patient compared to all WT samples. For HGPS-independent transcriptome changes we estimated average gene expression fold change in each WT compared to all other WT samples. **(C)** Heatmap of RNA-seq transcriptome analysis for DEGs consistently up- or down-regulated in 13 or more patients (80%). Heatmap of z-scaled, variance-stabilized gene expression values were generated using the Euclidean distance and Ward.D2 clustering method. All samples were analyzed in triplicate. **(D)** Heatmap of 50 GSEA hallmark pathways positively or negatively enriched in HGPS patients. Heatmap of mean-centered NES values was generated using Euclidean distance and complete linkage clustering method. **(E)** Gene Ontology Biological Processes (GOBP) positively (orange) or negatively (blue) enriched in the transcriptomes of HGPS patients. Relevant GOBP pathways were grouped into 15 representative biological functions, each containing 1-5 related pathways. Normalized Enrichment Score (NES) values from 16 HGPS patients for corresponding pathways were summarized as bar plots. Error bars represent the standard error of NES.

**Table 2.** Number of differentially expressed genes (DEGs) in each HGPS patient compared to all WT.

| **Patient** | **DEG**<br>**Up** | **DEG**<br>**Down** | **DEG**<br>**Total** |
| --- | --- | --- | --- |
| HGPS18 | 2795 | 3413 | 6208 |
| HGPS2 | 2386 | 2689 | 5075 |
| HGPS4 | 2266 | 2643 | 4909 |
| HGPS3 | 2043 | 2848 | 4891 |
| HGPS1 | 2182 | 2508 | 4690 |
| HGPS5 | 2009 | 2337 | 4346 |
| HGPS12 | 1951 | 2324 | 4275 |
| HGPS10 | 2338 | 1897 | 4235 |
| HGPS7 | 2053 | 2020 | 4073 |
| HGPS9 | 2373 | 1632 | 4005 |
| HGPS16 | 2048 | 1820 | 3868 |
| HGPS15 | 1787 | 2028 | 3815 |
| HGPS11 | 1591 | 1836 | 3427 |
| HGPS8 | 1513 | 1594 | 3107 |
| HGPS14 | 1584 | 1269 | 2853 |
| HGPS6 | 1709 | 1020 | 2729 |

To determine pathways that are consistently misregulated in HGPS patients, we performed Gene Set Enrichment Analysis (GSEA) and identified the top 19 pathways which are significantly (padj < 0.05) activated among the set of 50 GSEA hallmark pathways in HGPS patients (Fig. 2D; Table 3; Table S1)[43, 44]. Importantly, many of these pathways overlapped with those previously found to be misregulated based on studies in various cellular and mouse models of HGPS, including mTORC1 [26], Notch signaling [37], the UPR [29], Wnt/b-catenin signaling [28] and NF-kB[24], as well as pathways involved in myogenesis [45], oxidative phosphorylation [17] and apoptosis [46]. In addition, GSEA using the Gene Ontology (GO) pathway dataset highlighted biological processes involved in intermediate filament bundle assembly, cardiovascular and cardiac function, apoptosis, membrane invagination, ER stress pathways, cell adhesion and mitochondrial function to be significantly overrepresented in the set of up-regulated genes (Fig. 2E, Fig.S2A; Table S2), whereas biological processes involved in Wnt/b-catenin signaling, lipid metabolism and biosynthesis, adipogenesis, and retinoic acid receptor signaling pathway were significantly overrepresented in the set of down-regulated genes (Fig. 2E, Fig.S2A; Table S2). Again, several of these processes have previously been implicated in HGPS [6, 28, 29, 32, 36], validating our transcriptomics approach. Taken together, these data suggest that many of the major pathways that have been described to contribute to HGPS phenotypes in mouse and cellular disease models are also misregulated in progeria patients.

**Table 3.** 50 HALLMARK pathways in HGPS vs WT samples.

| Hallmark Pathway | pval | padj | log2err | ES | NES |
| --- | --- | --- | --- | --- | --- |
| HALLMARK_MTORC1_SIGNALING | 8.21E-09 | 2.71E-07 | 0.74774 | 0.495088 | 2.080073 |
| HALLMARK_EPITHELIAL_MESENCHYMAL_TRANSITION | 1.09E-08 | 2.71E-07 | 0.74774 | 0.483259 | 2.046614 |
| HALLMARK_NOTCH_SIGNALING | 0.005617 | 0.015604 | 0.407018 | 0.586773 | 1.842034 |
| HALLMARK_MYC_TARGETS_V2 | 0.001232 | 0.005134 | 0.45506 | 0.517382 | 1.787444 |
| HALLMARK_APICAL_JUNCTION | 3.45E-05 | 5.75E-04 | 0.557332 | 0.432323 | 1.78343 |
| HALLMARK_PI3K_AKT_MTOR_SIGNALING | 4.85E-04 | 0.003034 | 0.498493 | 0.47038 | 1.761829 |
| HALLMARK_GLYCOLYSIS | 5.12E-05 | 6.40E-04 | 0.557332 | 0.424201 | 1.75302 |
| HALLMARK_MYOGENESIS | 1.25E-04 | 0.001253 | 0.518848 | 0.418242 | 1.724709 |
| HALLMARK_UNFOLDED_PROTEIN_RESPONSE | 0.00157 | 0.005606 | 0.45506 | 0.432786 | 1.664837 |
| HALLMARK_HEDGEHOG_SIGNALING | 0.031702 | 0.066045 | 0.321776 | 0.531724 | 1.654135 |
| HALLMARK_WNT_BETA_CATENIN_SIGNALING | 0.02023 | 0.043978 | 0.352488 | 0.496132 | 1.628829 |
| HALLMARK_HYPOXIA | 6.86E-04 | 0.003812 | 0.477271 | 0.389109 | 1.625398 |
| HALLMARK_MYC_TARGETS_V1 | 3.54E-04 | 0.002528 | 0.498493 | 0.383753 | 1.624563 |
| HALLMARK_OXIDATIVE_PHOSPHORYLATION | 3.07E-04 | 0.002528 | 0.498493 | 0.383734 | 1.622497 |
| HALLMARK_ALLOGRAFT_REJECTION | 0.001208 | 0.005134 | 0.45506 | 0.411873 | 1.617142 |
| HALLMARK_TNFA_SIGNALING_VIA_NFKB | 8.23E-04 | 0.004115 | 0.477271 | 0.377455 | 1.573014 |
| HALLMARK_UV_RESPONSE_UP | 0.005491 | 0.015604 | 0.407018 | 0.379074 | 1.540065 |
| HALLMARK_ESTROGEN_RESPONSE_EARLY | 0.002565 | 0.008503 | 0.431708 | 0.360915 | 1.504083 |
| HALLMARK_MITOTIC_SPINDLE | 0.002721 | 0.008503 | 0.431708 | 0.351645 | 1.486819 |
| HALLMARK_KRAS_SIGNALING_DN | 0.054125 | 0.104086 | 0.321776 | 0.366282 | 1.395726 |
| HALLMARK_APOPTOSIS | 0.017847 | 0.040562 | 0.352488 | 0.342904 | 1.391452 |
| HALLMARK_ANGIOGENESIS | 0.075157 | 0.125261 | 0.237794 | 0.43533 | 1.354264 |
| HALLMARK_KRAS_SIGNALING_UP | 0.048802 | 0.097604 | 0.321776 | 0.322613 | 1.324355 |
| HALLMARK_INFLAMMATORY_RESPONSE | 0.063918 | 0.114138 | 0.257206 | 0.308445 | 1.253881 |
| HALLMARK_ADIPOGENESIS | 0.0625 | 0.114138 | 0.261664 | 0.290365 | 1.214904 |
| HALLMARK_IL2_STAT5_SIGNALING | 0.077869 | 0.125595 | 0.231127 | 0.291353 | 1.199337 |
| HALLMARK_P53_PATHWAY | 0.085239 | 0.133186 | 0.222056 | 0.283058 | 1.183051 |
| HALLMARK_CHOLESTEROL_HOMEOSTASIS | 0.187905 | 0.260979 | 0.148262 | 0.329372 | 1.172772 |
| HALLMARK_REACTIVE_OXYGEN_SPECIES_PATHWAY | 0.207627 | 0.27195 | 0.138805 | 0.341592 | 1.169288 |
| HALLMARK_ESTROGEN_RESPONSE_LATE | 0.2 | 0.27027 | 0.1396 | 0.269913 | 1.11043 |
| HALLMARK_E2F_TARGETS | 0.348269 | 0.404964 | 0.100633 | 0.247582 | 1.046821 |
| HALLMARK_G2M_CHECKPOINT | 0.347475 | 0.404964 | 0.100279 | 0.246001 | 1.038581 |
| HALLMARK_ANDROGEN_RESPONSE | 0.386892 | 0.43965 | 0.096563 | 0.275592 | 1.021037 |
| HALLMARK_DNA_REPAIR | 0.985626 | 0.985626 | 0.047543 | 0.185092 | 0.752738 |
| HALLMARK_TGF_BETA_SIGNALING | 0.932914 | 0.951953 | 0.051011 | 0.213552 | 0.735022 |
| HALLMARK_APICAL_SURFACE | 0.838403 | 0.873337 | 0.051426 | -0.2419 | -0.75179 |
| HALLMARK_INTERFERON_GAMMA_RESPONSE | 0.524085 | 0.557537 | 0.074739 | -0.23822 | -0.98035 |
| HALLMARK_PROTEIN_SECRETION | 0.451796 | 0.491082 | 0.081527 | -0.26801 | -1.00226 |
| HALLMARK_SPERMATOGENESIS | 0.422053 | 0.468948 | 0.085536 | -0.27844 | -1.01902 |
| HALLMARK_PEROXISOME | 0.292776 | 0.357043 | 0.10714 | -0.29065 | -1.07829 |
| HALLMARK_UV_RESPONSE_DN | 0.212121 | 0.27195 | 0.128789 | -0.28215 | -1.12741 |
| HALLMARK_FATTY_ACID_METABOLISM | 0.218519 | 0.273148 | 0.125033 | -0.28672 | -1.13653 |
| HALLMARK_HEME_METABOLISM | 0.158607 | 0.226582 | 0.153159 | -0.28688 | -1.1669 |
| HALLMARK_INTERFERON_ALPHA_RESPONSE | 0.138196 | 0.203229 | 0.164406 | -0.32315 | -1.20375 |
| HALLMARK_COAGULATION | 0.071429 | 0.123153 | 0.234393 | -0.34576 | -1.29064 |
| HALLMARK_PANCREAS_BETA_CELLS | 0.097328 | 0.147467 | 0.197822 | -0.51325 | -1.39423 |
| HALLMARK_COMPLEMENT | 0.014756 | 0.035134 | 0.38073 | -0.34578 | -1.39458 |
| HALLMARK_XENOBIOTIC_METABOLISM | 0.010993 | 0.028929 | 0.38073 | -0.34527 | -1.39868 |
| HALLMARK_BILE_ACID_METABOLISM | 0.014222 | 0.035134 | 0.38073 | -0.41186 | -1.52797 |
| HALLMARK_IL6_JAK_STAT3_SIGNALING | 0.001522 | 0.005606 | 0.45506 | -0.49147 | -1.7326 |

### HGPS patients show uniform deregulation of pathways

The observed wide range of clinical presentation and different rate of progression of HGPS patients [8, 9] raises the question of whether the same pathways are pervasively misregulated across the patient population or whether the set of misregulated genes varies between patients. To address this question, we compared the behavior of several of the major misregulated pathways including mTORC1 signaling, Notch and mesodermal cell specification pathways in the set of HGPS patients. We generated enrichment score (ES) plots for each HGPS patient (n=16; red lines) and each WT control (n=10; green lines) (Fig. 3A-C). Misregulation of these pathways was consistently observed across most patients and the direction of normalized enrichment scores (NES), either positive or negative, for the three pathways examined was also shared by the majority of patients (Fig.3A-C). For example, 15 of 16 patients showed activation of mTORC1 signaling (Fig. 3A), all patients exhibited enhanced Notch signaling (Fig. 3B), and 13 of 16 patients presented with downregulation of mesodermal cell fate programs, while one showed slight upregulation and two were unaffected (Fig. 3C). To generalize this analysis, we extended it to the top ten positively and negatively enriched pathways in all HGPS and WT samples (Figure 3D). The distribution of NES values across all HGPS samples suggested limited heterogeneity in the patient population (Fig. 3D, E). All patients showed enrichment of at least 28 of the 50 pathways, and 50% of HGPS patients showed positive enrichment of at least 35 pathways (NES > 0), (Fig. 3 E). Importantly, the most prominent HGPS pathways, including mTORC1, Notch signaling and the UPR, as well as pathways involved in myogenesis, oxidative phosphorylation, apoptosis, inflammatory response, ROS production, DNA repair and angiogenesis were upregulated in 80-100% of HGPS patients (Fig. 3E, Fig. S2B). Taken together, our data suggests that HGPS patients exhibit relatively uniform misregulation of core disease-related pathways, especially of the key pathways affected in HGPS.

**Figure 3.**
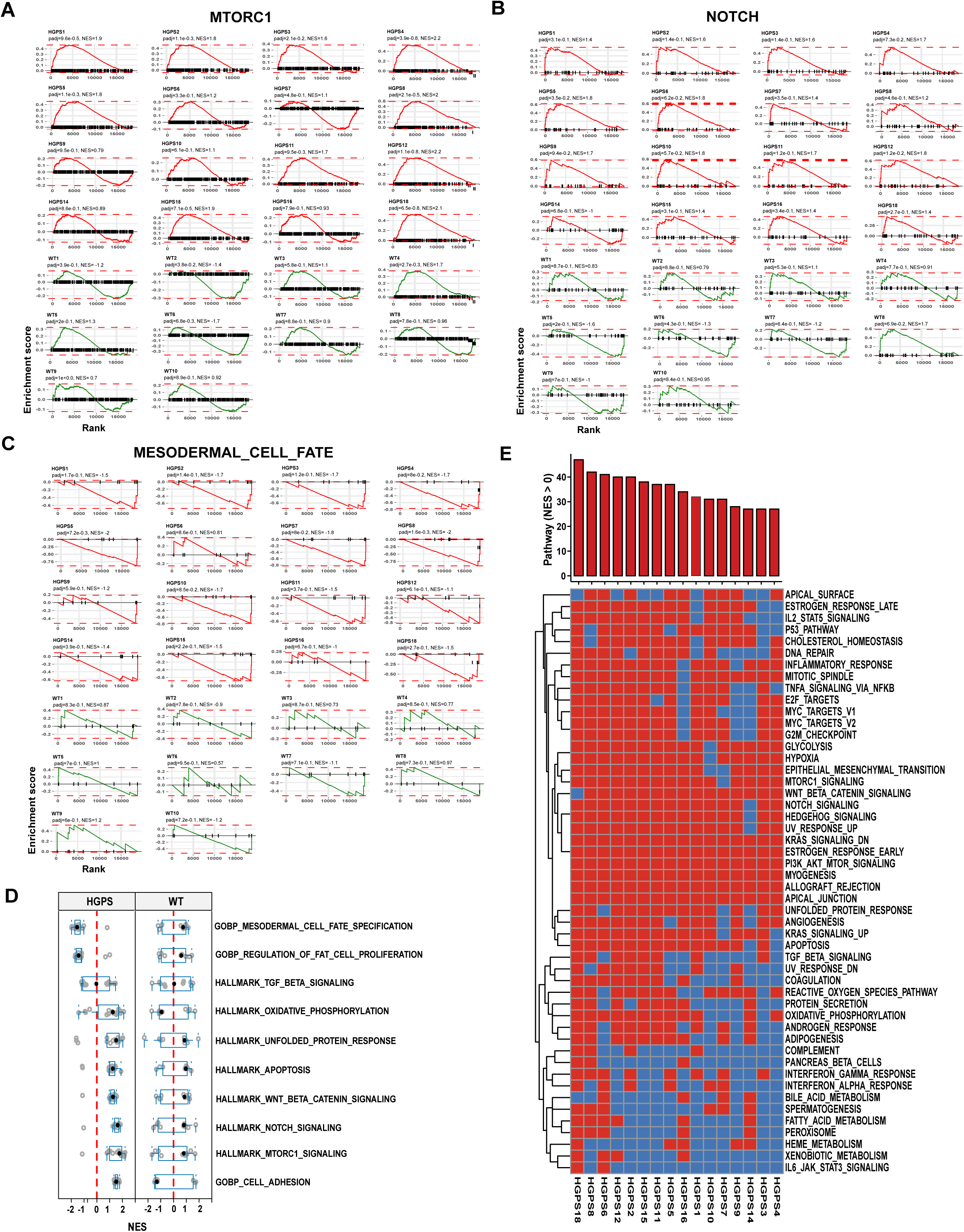
Pathway deregulation in HGPS patients. **(A-C)** Enrichment score plots for two positively enriched pathways **(A, B)** and one negatively enriched pathway **(C)** in 16 HGPS patients (red) and 10 WT controls (green). Red or green lines represent the running enrichment score (ES) calculated from the ranked gene list (Wald statistic) for each dataset. The peaks near the top left or bottom right indicate pathway alterations driven by subsets of up- or down-regulated genes. Adjusted p values (padj) and normalized enrichment scores (NES) are shown for each patient. **(D)** Boxplots of 10 positively or negatively enriched pathways in HGPS shown as NES values across all 16 HGPS patients and 10 WT controls. Individual datapoints represent the normalized enrichment score (NES) for each patient for the given pathway. Boxes represent the interquartile range (25th to 75th percentiles), and the whiskers extend to the 5th and 95th percentiles. **(E)** Binary representation of 50 GSEA hallmark pathways in individual HGPS patients based on NES. Red represents positively enriched pathways (NES > 0) and blue represents negatively enriched pathways (NES < 0). Top bar chart represents the number of pathways that are positively enriched in a single patient.

### Comparison between a progerin-inducible cell line and primary patient fibroblasts identifies immediate targets of progerin

A commonly used cellular model to probe HGPS mechanisms are immortalized skin fibroblasts which ectopically express progerin [40, 47–49]. To understand how well HGPS cellular models recapitulate the disease, we compared the transcriptional status of all 16 primary patient skin fibroblast samples to a well-characterized GFP-progerin inducible skin fibroblast cell line after 6 days of induction, which recapitulates many of the most prominent hallmarks of HGPS patient cells, including altered nuclear morphology, epigenetic marks and increased DNA damage [40] (Fig. S3A). This approach allows us to identify immediate early pathways in HGPS which are directly triggered by induction of progerin and to distinguish them from adaptive organismal pathways which may be activated in the primary patient cells during prolonged progerin exposure.

DGE analysis of uninduced control cells and GFP-progerin expressing cells identified a total of 4583 differentially expressed genes, of which 2297 were upregulated and 2286 were downregulated in GFP-progerin expressing cells (Fig. 4A; FDR cut off: 0.05, LFC cut off: log2(1.5); Fig. S3A). These DEGs were compared with the 693 genes consistently up- or down-regulated in HGPS patient samples (Fig.2 C) and we identified a total of 148 differentially expressed genes between all primary cell lines and GFP-progerin induced cells, of which 85 genes were upregulated and 63 downregulated in both GFP-progerin and HGPS patient cell lines (Fig. 4B).

**Figure 4.**
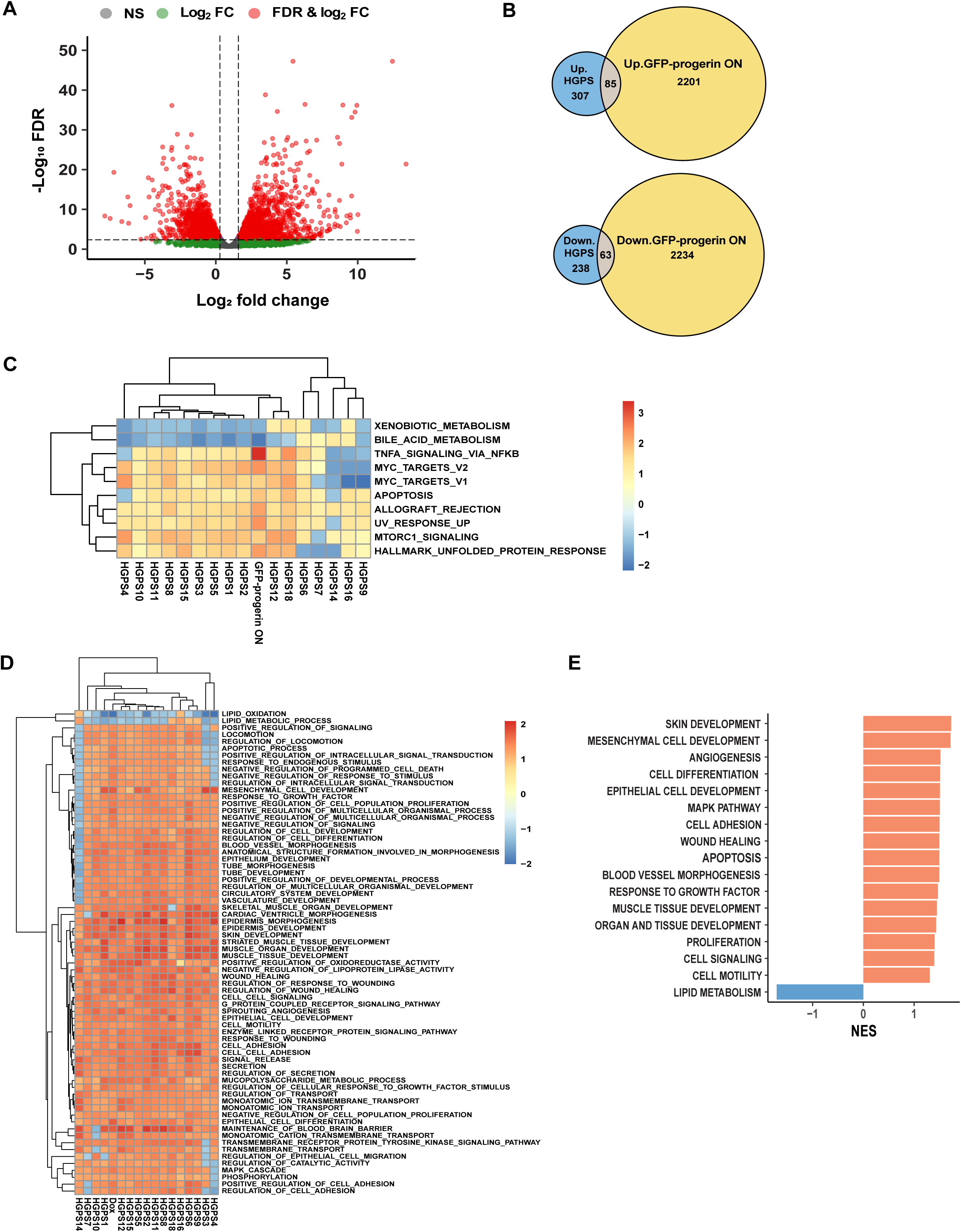
Pathway comparison between primary HGPS patient fibroblasts and GFP-progerin inducible fibroblasts. **(A)** Volcano plot comparison of differentially expressed genes upon induced expression of GFP-progerin for 6 days and uninduced control cells. Dashed lines indicate threshold (FDR < 0.05, |log2FC| > log2(1.5)). Data point colors denote whether the criteria for FC, FDR, or both were met. **(B)** Up-regulated (upper panel) and down-regulated (lower panel) DEGs in primary HGPS fibroblasts (blue), GFP-progerin expressing cells (yellow) and deregulated in both (brown). **(C)** Heatmap of the top 10 GSEA hallmark pathways significantly affected (padj<0.05) with consistent positive or negative trends in both HGPS patients and GFP-progerin expressing cells. **(D)** Heatmap of the top 70 Gene Ontology biological processes (GOBP) GSEA pathways significantly affected (padj<0.05) with consistent positive or negative trends in both HGPS patients and GFP-progerin expressing cells. Heatmap of NES values was generated using Euclidean distance and complete linkage clustering method. **(E)** GOBP GSEA pathways showing similar positive (red) or negative (blue) enrichment trends in the transcriptomes of both HGPS patients and GFP-progerin expressing cells.

To determine which pathways are distinct and which ones are shared in the HGPS cell model and HGPS patients, we compared pathway alterations measured as NES from GSEA using the 50 hallmark gene sets in the inducible cell line, all HGPS cell lines, and all WT cell lines. PCA based on NES values of the 50 hallmark pathways identified in the patients showed that the GFP-progerin-expressing cell line clustered more closely with HGPS patient cells than WT samples, suggesting that the affected pathways in the progerin cell line are more similar to HGPS than to the WT samples (Fig. S3C). Of the 19 hallmark GSEA pathways identified in patient samples, nine were also significantly affected in GFP-progerin expressing cells (Fig.4C and S3D; Table 4, magenta; Table S3) including mTORC1, the UV response, apoptosis, the UPR and TNFα signaling via NF-κB. Thirteen pathways were significantly affected in GFP-progerin expressing cells only, including the inflammatory response, KRAS and p53 signaling, G2/M checkpoint and adipogenesis (Table 4, green; Table S3), whereas eight pathways were significantly misregulated only in HGPS patients, including oxidative phosphorylation, Notch signaling and Wnt/b-catenin signaling (Table 4, red; Table S3), possibly reflecting adaptive responses in patients. Interestingly, six pathways were found to be significantly affected in both GFP-progerin expressing cells and HGPS patients but in opposite directions (Table 4, blue). For example, myogenesis and epithelial to mesenchymal transition were found to be repressed in GFP-progerin cells but activated in HGPS patients, suggesting that these pathways are a result of the physiological response to progerin in HGPS patients that is absent in cell culture conditions. In addition, GSEA using the Gene Ontology (GO) pathway dataset highlighted biological processes involved in cell adhesion, cell signaling, apoptosis, organ and tissue development, cell differentiation and proliferation, angiogenesis and blood vessel morphogenesis as significantly overrepresented in the set of HGPS-dependent up-regulated genes in both the inducible cell line and all HGPS cell lines (Fig. 4D, E, Table S4), whereas biological processes involved in lipid metabolism were significantly overrepresented in the set of HGPS-dependent down-regulated genes in both the inducible cell line and all HGPS cell lines (Fig. 4D, E, Table S4).

**Table 4.** 50 HALLMARK pathways in GFP-progerin cell line and HGPS samples. Table is sorted by the NES values of the GFP-progerin cell line. Pathways similarly affected in both cell lines are shown in magenta, pathways affected only in GFP-progerin cell line are shown in green and pathways affected only in HGPS cells are shown in red. Blue color represents pathways that are significantly affected in both cell lines but show a different direction. Black color represents pathways that are not significantly affected.

| Hallmark Pathway | GFP-progerin padj | GFP-progerin NES | HGPS padj | HGPS NES |
| --- | --- | --- | --- | --- |
| HALLMARK_TNFA_SIGNALING_VIA_NFKB | 2.35E-42 | 3.37 | 4.11E-03 | 1.57 |
| HALLMARK_INFLAMMATORY_RESPONSE | 3.19E-23 | 2.94 | 1.14E-01 | 1.25 |
| HALLMARK_INTERFERON_ALPHA_RESPONSE | 6.53E-15 | 2.77 | 2.03E-01 | -1.20 |
| HALLMARK_INTERFERON_GAMMA_RESPONSE | 4.76E-17 | 2.64 | 5.58E-01 | -0.98 |
| HALLMARK_UV_RESPONSE_UP | 5.85E-10 | 2.33 | 1.56E-02 | 1.54 |
| HALLMARK_MYC_TARGETS_V2 | 7.09E-07 | 2.33 | 5.13E-03 | 1.79 |
| HALLMARK_KRAS_SIGNALING_UP | 2.45E-10 | 2.28 | 9.76E-02 | 1.32 |
| HALLMARK_UNFOLDED_PROTEIN_RESPONSE | 4.08E-08 | 2.24 | 5.61E-03 | 1.66 |
| HALLMARK_ALLOGRAFT_REJECTION | 1.19E-07 | 2.17 | 5.13E-03 | 1.62 |
| HALLMARK_COMPLEMENT | 4.71E-06 | 1.96 | 3.51E-02 | -1.39 |
| HALLMARK_IL6_JAK_STAT3_SIGNALING | 1.86E-03 | 1.78 | 5.61E-03 | -1.73 |
| HALLMARK_APOPTOSIS | 2.31E-04 | 1.76 | 4.06E-02 | 1.39 |
| HALLMARK_P53_PATHWAY | 1.43E-04 | 1.75 | 1.33E-01 | 1.18 |
| HALLMARK_IL2_STAT5_SIGNALING | 3.84E-04 | 1.70 | 1.26E-01 | 1.20 |
| HALLMARK_MTORC1_SIGNALING | 2.31E-04 | 1.69 | 2.71E-07 | 2.08 |
| HALLMARK_G2M_CHECKPOINT | 5.23E-04 | 1.65 | 4.05E-01 | 1.04 |
| HALLMARK_MYC_TARGETS_V1 | 7.51E-04 | 1.62 | 2.53E-03 | 1.62 |
| HALLMARK_E2F_TARGETS | 1.29E-03 | 1.59 | 4.05E-01 | 1.05 |
| HALLMARK_WNT_BETA_CATENIN_SIGNALING | 1.99E-01 | 1.28 | 4.40E-02 | 1.63 |
| HALLMARK_ESTROGEN_RESPONSE_EARLY | 1.15E-01 | 1.23 | 8.50E-03 | 1.50 |
| HALLMARK_HYPOXIA | 1.48E-01 | 1.19 | 3.81E-03 | 1.63 |
| HALLMARK_ANDROGEN_RESPONSE | 2.34E-01 | 1.15 | 4.40E-01 | 1.02 |
| HALLMARK_ESTROGEN_RESPONSE_LATE | 2.34E-01 | 1.14 | 2.70E-01 | 1.11 |
| HALLMARK_TGF_BETA_SIGNALING | 3.13E-01 | 1.12 | 9.52E-01 | 0.74 |
| HALLMARK_DNA_REPAIR | 4.01E-01 | 1.04 | 9.86E-01 | 0.75 |
| HALLMARK_PI3K_AKT_MTOR_SIGNALING | 4.34E-01 | 1.04 | 3.03E-03 | 1.76 |
| HALLMARK_CHOLESTEROL_HOMEOSTASIS | 5.04E-01 | 0.99 | 2.61E-01 | 1.17 |
| HALLMARK_SPERMATOGENESIS | 8.39E-01 | 0.83 | 4.69E-01 | -1.02 |
| HALLMARK_NOTCH_SIGNALING | 8.39E-01 | 0.76 | 1.56E-02 | 1.84 |
| HALLMARK_PROTEIN_SECRETION | 9.91E-01 | 0.66 | 4.91E-01 | -1.00 |
| HALLMARK_PANCREAS_BETA_CELLS | 7.80E-01 | -0.82 | 1.47E-01 | -1.39 |
| HALLMARK_GLYCOLYSIS | 3.87E-01 | -1.04 | 6.40E-04 | 1.75 |
| HALLMARK_ANGIOGENESIS | 3.63E-01 | -1.11 | 1.25E-01 | 1.35 |
| HALLMARK_REACTIVE_OXYGEN_SPECIES_PATHWAY | 2.65E-01 | -1.15 | 2.72E-01 | 1.17 |
| HALLMARK_HEME_METABOLISM | 1.54E-01 | -1.21 | 2.27E-01 | -1.17 |
| HALLMARK_KRAS_SIGNALING_DN | 1.24E-01 | -1.27 | 1.04E-01 | 1.40 |
| HALLMARK_MITOTIC_SPINDLE | 3.65E-02 | -1.33 | 8.50E-03 | 1.49 |
| HALLMARK_XENOBIOTIC_METABOLISM | 2.86E-02 | -1.36 | 2.89E-02 | -1.40 |
| HALLMARK_HEDGEHOG_SIGNALING | 1.16E-01 | -1.39 | 6.60E-02 | 1.65 |
| HALLMARK_APICAL_JUNCTION | 2.86E-02 | -1.43 | 5.75E-04 | 1.78 |
| HALLMARK_COAGULATION | 2.71E-02 | -1.46 | 1.23E-01 | -1.29 |
| HALLMARK_ADIPOGENESIS | 2.94E-03 | -1.50 | 1.14E-01 | 1.21 |
| HALLMARK_APICAL_SURFACE | 5.58E-02 | -1.52 | 8.73E-01 | -0.75 |
| HALLMARK_OXIDATIVE_PHOSPHORYLATION | 1.25E-03 | -1.56 | 2.53E-03 | 1.62 |
| HALLMARK_UV_RESPONSE_DN | 2.21E-03 | -1.59 | 2.72E-01 | -1.13 |
| HALLMARK_MYOGENESIS | 1.25E-03 | -1.63 | 1.25E-03 | 1.72 |
| HALLMARK_FATTY_ACID_METABOLISM | 5.75E-04 | -1.68 | 2.73E-01 | -1.14 |
| HALLMARK_EPITHELIAL_MESENCHYMAL_TRANSITION | 1.04E-04 | -1.72 | 2.71E-07 | 2.05 |
| HALLMARK_PEROXISOME | 2.08E-04 | -1.88 | 3.57E-01 | -1.08 |
| HALLMARK_BILE_ACID_METABOLISM | 1.87E-05 | -2.03 | 3.51E-02 | -1.53 |

Taken together, our data suggests mTORC1, the UV response, apoptosis, the UPR and TNFα signaling via NF-κB pathways as immediate targets of progerin, whereas oxidative phosphorylation, Notch signaling and Wnt/b-catenin signaling likely reflect adaptive or compensatory pathways to long-term progerin expression *in vivo* in HGPS patients.

## DISCUSSION

We have transcriptionally profiled a collection of HGPS patient samples. Our analysis has enabled the identification of several major cellular pathways which are dysregulated across HGPS patients. Our analysis also demonstrates that gene expression profiles are similar amongst patients despite documented heterogeneity of clinical symptoms in the HGPS patient population. Furthermore, by comparing patient transcriptome profiles with an inducible cell-based HGPS model, we identify primary pathways affected by the presence of the disease-causing progerin protein and distinguish them from secondary adaptive pathways that emerge in patients during disease progression.

Progeroid syndromes are a group of diseases characterized by rapid premature aging. One of the most prominent forms of progeria is HGPS [2], which is caused by a single point mutation (c.1824C>T; G608G) in exon 11 of the *LMNA* gene, resulting in the alternatively spliced mRNA and subsequent production of the mutant lamin A protein progerin [1, 2]. Other progeroid syndromes can be caused by mutations in *LMNA* other than c.1824C>T and are referred to as progeroid laminopathies or can be due to non-*LMNA* mutations which are classified as atypical progeroid syndromes [7]. In addition, there are patients with non-classic HGPS mutations in *LMNA* which lead to progerin production in the absence of the c.1824C>T mutation [50, 51]. The various types of progeria show overlapping organismal phenotypes which can make diagnosis and classification in the absence of genetic or molecular markers difficult [52–54]. This challenge is illustrated by our observation that four of twenty patient samples available in a public cell repository which are classified as HGPS patient samples based on patient phenotype lacked the classic HGPS c.1824C>T mutation in the *LMNA* gene and no progerin mRNA was detected. The distinct nature of these cell lines was further underscored by their segregation from classic HGPS samples in PCA of transcriptomics data, indicating that they do not represent classic HGPS patients. Our findings highlight that caution should be taken when using publicly available patient cell lines, especially when there is no available sequencing data confirming the presence of the classical HGPS mutation.

Transcriptomic profiling of patient samples confirmed the disease relevance of several cellular pathways which have previously been implicated in HGPS in cellular and mouse models of the disease [55, 56]. One prominent pathway identified by transcriptional profiling and previously implicated in HGPS is mTORC1 signaling. Genetic reduction of <u>m</u>echanistic target <u>o</u>f rapamycin (mTOR) significantly extends lifespan in HGPS mouse models, implicating hyperactivation of mTORC1 signaling in the acceleration of age-associated cellular decline [26, 27]. In addition, dysregulated mTOR activity likely contributes to impaired autophagy and metabolic imbalance, thereby exacerbating progerin-induced cellular stress [57, 58]. Transcriptional profiling also identified misregulation of pathways which contribute to endothelial-to-mesenchymal transition (EndMT) as shared in many patients. This finding is in line with the demonstration of a detrimental role of progerin in mesenchymnal stem cell function and differentiation [37]. In addition, vascular cell pathology is a major cause of mortality in HGPS [6]. Recent findings demonstrate that EndMT is a key contributor to accelerated atherosclerosis in HGPS mice, suggesting that vascular remodeling and fibrosis are driven by lineage reprogramming of endothelial cells under progerin stress [25]. Another prominent dysregulated signaling pathway shared amongst patients is the Notch signaling pathway. Activation of canonical Notch effectors in response to progerin expression has been observed in cultured cells, including mesenchymal stem cells [37]. Given the role of Notch in regulating cell fate and tissue homeostasis, such dysregulation may impair regenerative capacity and exacerbate the degenerative phenotype of HGPS. In parallel, we also find dysregulation of the cellular unfolded protein response (UPR), which has been implicated in promoting apoptosis and vascular dysfunction in progeria models, consistent with the finding that progerin-induced protein misfolding leads to chronic endoplasmic reticulum (ER) stress [18, 29]. Abnormalities of the nuclear lamina in HGPS have also previously been reported to trigger constitutive activation of NF-κB signaling, resulting in a systemic pro-inflammatory state [24]. In agreement with these findings, we observe prevalent dysregulation of NF-kB signaling in most patient cells. Chronic inflammation is a hallmark of both HGPS and normal aging and likely serves as a major driver of tissue dysfunction and degeneration [38, 59]. Similarly, we find consistent upregulation of oxidative stress response pathways in patients. Oxidative stress represents another key pathogenic mechanism in HGPS, as impaired NRF2 activity or increased reactive oxygen species (ROS) levels are sufficient to recapitulate HGPS-associated phenotypes [17, 32, 60]. Collectively, these findings underscore the multifactorial nature of HGPS pathogenesis, implicating interconnected signaling cascades involved in inflammation, oxidative stress, proteostasis, and vascular remodeling. Reassuringly, our findings indicate that many of the major pathways that have been described to contribute to HGPS phenotypes in mouse and cellular disease models are also misregulated in progeria patients and targeting these pathways may provide therapeutic avenues to mitigate disease severity and improve outcomes in HGPS.

Although individuals with HGPS typically exhibit a characteristic set of clinical features, such as craniofacial abnormalities, growth retardation, and cardiovascular complications, there is notable variability in the age of onset, severity, and progression of symptoms between patients [7, 9]. At the cellular level, HGPS is associated with several hallmark abnormalities, including nuclear envelope defects, decreased expression of several nuclear proteins and epigenetic marks, mitochondrial dysfunction, and increased cellular senescence [1, 11, 30, 61]. These cellular phenotypes also exhibit considerable variation between patients, possibly contributing to differences in clinical outcomes. Our results indicate that even though some degree of transcriptional heterogeneity between the individual patients exists, the majority of patients exhibit misregulation of a set of shared pathways, suggesting that these pathways are universal driver mechanisms in HGPS. Further research is needed to understand the molecular and genetic factors that underlie inter-individual variability in disease expression and progression.

A limitation of pathway analysis of HGPS patient samples is to distinguish the pathways which are directly targeted by the disease-causing progerin protein and the emergence of adaptive secondary response pathways during progression of the disease in patients during their lifetime. The same caveat applies to the use of cell-based models used in the study of HGPS disease mechanisms. To identify pathways which are directly triggered by induction of progerin, we took advantage of the availability of immortalized skin fibroblasts expressing GFP-tagged progerin in an inducible manner, which have served as a powerful cellular model commonly used to probe HGPS mechanisms [17, 18, 40, 62]. We compared the set of cellular pathways dysregulated in HGPS patients to those altered by acute expression of progerin. GSEA analysis identified nine hallmark pathways significantly affected in both primary HGPS fibroblasts and GFP-progerin expressing cells, including mTORC1, the UV response, apoptosis, the UPR and TNFα signaling via NF-κB. We consider these pathways disease-relevant immediate targets of the disease-causing progerin protein. In contrast, other pathways such as oxidative phosphorylation, Notch signaling and Wnt/b-catenin signaling were only seen misregulated in patient cells but not upon acute progerin expression, suggesting that they arise as an adaptive or compensatory response during chronic disease progression. We also identified the Myc pathway as well as allograft rejection pathway as significantly upregulated in direct response to progerin and in patients. Although myc expression is generally associated with cancer, HGPS patients do not have an increased incidence of cancer, possibly due to an inherent resistance mechanism mediated by the BRD4 transcription factor [63]. Myc may, however, contribute to the premature aging process in HGPS rather than tumor formation. While allograft rejection may seem a surprising progerin-target pathway, many of the genes attributed to this pathway are involved in inflammation, which is well documented in HGPS [38].

Taken together, our analysis provides a systematic, albeit partial, transcriptional profile of HGPS patients. The results reassuringly confirm the involvement of various previously implicated major cellular pathways in HGPS, and they also point to some novel aspects of HGPS. We hope this collection of transcriptional profiles will serve as a foundation for the continued elucidation of disease mechanisms and development of therapeutics for this rare disease.

## AUTHOR CONTRIBUTIONS

S.V. and T.M. designed the study and wrote the manuscript; S.V. performed RNA-seq experiments and S.K. performed RNA-seq analysis. All authors read and approved the manuscript.

## ACKNOWLEDGMENTS

We thank the Misteli lab members for sharing feedback, data, and reagents. We are grateful to Leslie Gordon and Progeria Research Foundation for providing us with primary HGPS patient fibroblasts and for comments on the manuscript. RNA-seq was conducted at the Center for Cancer Research (CCR) Genomics Core and CCR Sequencing Facility at the NCI. This research was supported by the Intramural Research Program of the NIH, NCI, Center for Cancer Research through grant 1-ZIA-BC010309-25 to T.M. The contributions of the NIH author(s) were made as part of their official duties as NIH federal employees, are in compliance with agency policy requirements, and are considered Works of the United States Government. However, the findings and conclusions presented in this paper are those of the author(s) and do not necessarily reflect the views of the NIH or the U.S. Department of Health and Human Services.

