## Supplemental Figures for "Transcriptional profiling of Hutchinson-Gilford Progeria patients identifies primary target pathways of progerin"

### Figure S1

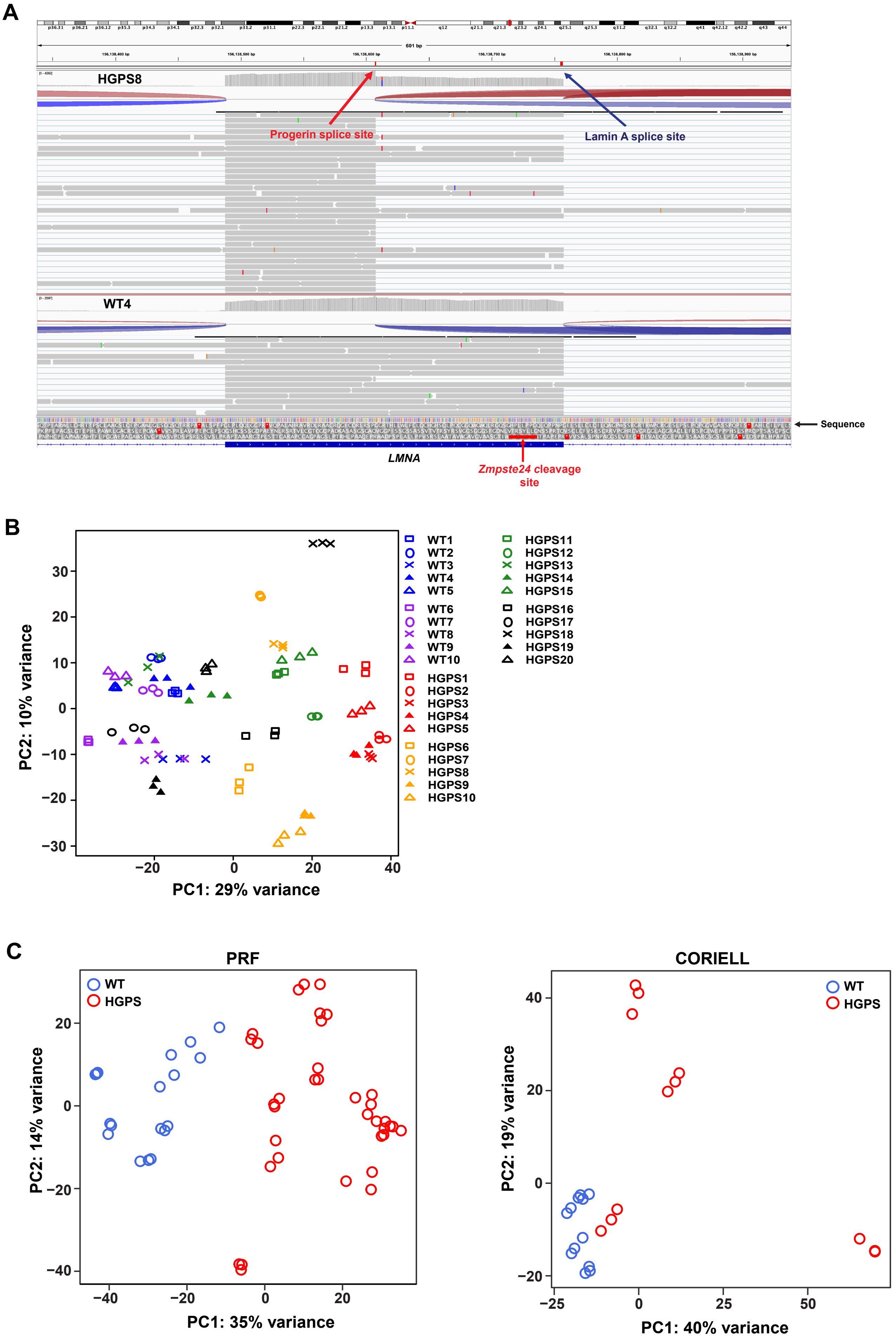
A
HGPS8
Progerin splice site
Lamin A splice site
WT4
Sequence
LMNA
Zmpste24 cleavage site
B
PC1: 29% variance
PC2: 10% variance
WT1
WT2
WT3
WT4
WT5
WT6
WT7
WT8
WT9
WT10
HGPS1
HGPS2
HGPS3
HGPS4
HGPS5
HGPS6
HGPS7
HGPS8
HGPS9
HGPS10
HGPS11
HGPS12
HGPS13
HGPS14
HGPS15
HGPS16
HGPS17
HGPS18
HGPS19
HGPS20
C
PRF
WT
HGPS
PC1: 35% variance
PC2: 14% variance
CORIELL
PC1: 40% variance
PC2: 19% variance

#### SUPPLEMENTAL FIGURE LEGENDS

**Figure S1. Characterization of primary human dermal fibroblasts. (A)** Integrative Genomics Viewer (IGV) screen capture showing the two splicing junctions at chr1:156,138,607 (progerin) and chr: 156,138,757 (Lamin A). Sequence reads at these sites were used to quantitate progerin and wild type lamin A transcripts in a control and HGPS patient. **(B)** Principal component analysis (PCA) of the primary fibroblast datasets used in this study. Samples primarily segregate by progeria/control phenotype with four samples classified by the NIA Aging Cell Repository (Coriell) as progeria patients showing relatively similar gene expression patterns as the WT samples. Follow-up analysis demonstrated absence of the classic c.1824C>T mutation in these samples and identified them as atypical progerias. These samples were excluded from analysis. Top 500 genes with the highest variability across samples were used for PCA plot. All samples were analyzed in triplicate. **(C)** PCA of the primary fibroblast datasets based on the source (PRF Cell and Tissue Bank, Coriell NIA Aging Cell Repository). Samples primarily segregate by progeria/control phenotype. Blue dots represent wild type control samples and red dots represent progeria patients. All samples were analyzed in triplicate.

### Figure S2

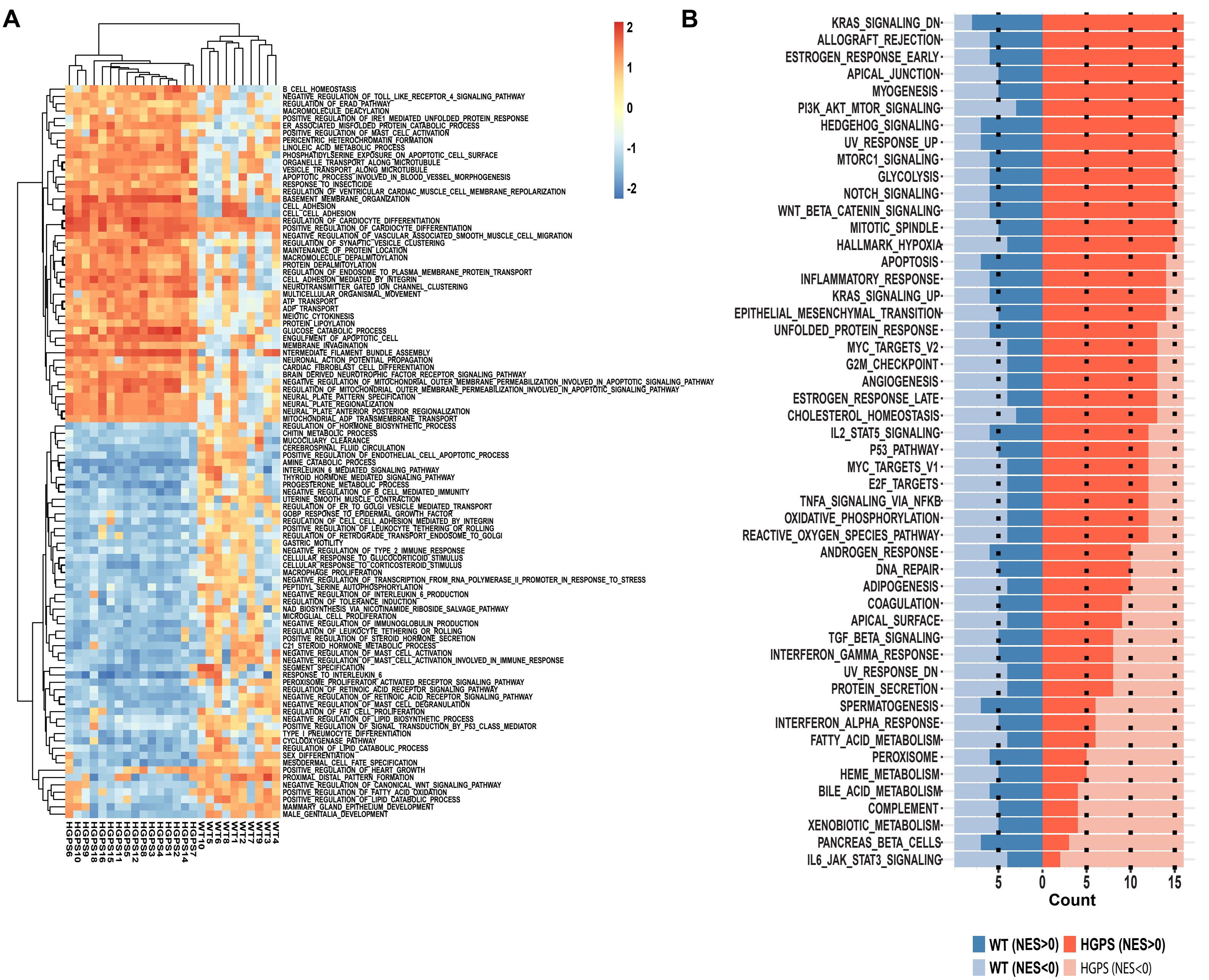
A
2
1
0
-1
-2
B_CELL_HOMEOSTASIS
NEGATIVE_REGULATION_OF_TOLL_LIKE_RECEPTOR_4_SIGNALING_PATHWAY
REGULATION_OF_ERAD_PATHWAY
MACROMOLECULE_DEACYLATION
POSITIVE_REGULATION_OF_IRE1_MEDIATED_UNFOLDED_PROTEIN_RESPONSE
ER_ASSOCIATED_MISFOLDED_PROTEIN_CATABOLIC_PROCESS
POSITIVE_REGULATION_OF_MAST_CELL_ACTIVATION
PERICENTRIC_HETEROCHROMATIN_FORMATION
LINOLEIC_ACID_METABOLIC_PROCESS
PHOSPHATIDYLSERINE_EXPOSURE_ON_APOPTOTIC_CELL_SURFACE
ORGANELLE_TRANSPORT_ALONG_MICROTUBULE
VESICLE_TRANSPORT_ALONG_MICROTUBULE
APOPTOTIC_PROCESS_INVOLVED_IN_BLOOD_VESSEL_MORPHOGENESIS
RESPONSE_TO_INSECTICIDE
REGULATION_OF_VENTRICULAR_CARDIAC_MUSCLE_CELL_MEMBRANE_REPOLARIZATION
BASEMENT_MEMBRANE_ORGANIZATION
CELL_ADHESION
CELL_CELL_ADHESION
REGULATION_OF_CARDIOCYTE_DIFFERENTIATION
POSITIVE_REGULATION_OF_CARDIOCYTE_DIFFERENTIATION
NEGATIVE_REGULATION_OF_VASCULAR_ASSOCIATED_SMOOTH_MUSCLE_CELL_MIGRATION
REGULATION_OF_SYNAPTIC_VESICLE_CLUSTERING
MAINTENANCE_OF_PROTEIN_LOCATION
MACROMOLECULE_DEPALMITOYLATION
PROTEIN_DEPALMITOYLATION
REGULATION_OF_ENDOSOME_TO_PLASMA_MEMBRANE_PROTEIN_TRANSPORT
CELL_ADHESION_MEDIATED_BY_INTEGRIN
NEUROTRANSMITTER_GATED_ION_CHANNEL_CLUSTERING
MULTICELLULAR_ORGANISMAL_MOVEMENT
ATP_TRANSPORT
ADP_TRANSPORT
MEIOTIC_CYTOKINESIS
PROTEIN_LIPOYLATION
GLUCOSE_CATABOLIC_PROCESS
ENGULFMENT_OF_APOPTOTIC_CELL
MEMBRANE_INVAGINATION
INTERMEDIATE_FILAMENT_BUNDLE_ASSEMBLY
NEURONAL_ACTION_POTENTIAL_PROPAGATION
CARDIAC_FIBROBLAST_CELL_DIFFERENTIATION
BRAIN_DERIVED_NEUROTROPHIC_FACTOR_RECEPTOR_SIGNALING_PATHWAY
NEGATIVE_REGULATION_OF_MITOCHONDRIAL_OUTER_MEMBRANE_PERMEABILIZATION_INVOLVED_IN_APOPTOTIC_SIGNALING_PATHWAY
REGULATION_OF_MITOCHONDRIAL_OUTER_MEMBRANE_PERMEABILIZATION_INVOLVED_IN_APOPTOTIC_SIGNALING_PATHWAY
NEURAL_PLATE_PATTERN_SPECIFICATION
NEURAL_PLATE_REGIONALIZATION
NEURAL_PLATE_ANTERIOR_POSTERIOR_REGIONALIZATION
MITOCHONDRIAL_ADP_TRANSMEMBRANE_TRANSPORT
REGULATION_OF_HORMONE_BIOSYNTHETIC_PROCESS
CHITIN_METABOLIC_PROCESS
MUCOCILIARY_CLEARANCE
CEREBROSPINAL_FLUID_CIRCULATION
POSITIVE_REGULATION_OF_ENDOTHELIAL_CELL_APOPTOTIC_PROCESS
AMINE_CATABOLIC_PROCESS
INTERLEUKIN_6_MEDIATED_SIGNALING_PATHWAY
THYROID_HORMONE_MEDIATED_SIGNALING_PATHWAY
PROGESTERONE_METABOLIC_PROCESS
NEGATIVE_REGULATION_OF_B_CELL_MEDIATED_IMMUNITY
UTERINE_SMOOTH_MUSCLE_CONTRACTION
REGULATION_OF_ER_TO_GOLGI_VESICLE_MEDIATED_TRANSPORT
GOBP_RESPONSE_TO_EPIDERMAL_GROWTH_FACTOR
REGULATION_OF_CELL_CELL_ADHESION_MEDIATED_BY_INTEGRIN
POSITIVE_REGULATION_OF_LEUKOCYTE_TETHERING_OR_ROLLING
REGULATION_OF_RETROGRADE_TRANSPORT_ENDOSOME_TO_GOLGI
GASTRIC_MOTILITY
NEGATIVE_REGULATION_OF_TYPE_2_IMMUNE_RESPONSE
CELLULAR_RESPONSE_TO_GLUCOCORTICOID_STIMULUS
CELLULAR_RESPONSE_TO_CORTICOSTEROID_STIMULUS
MACROPHAGE_PROLIFERATION
NEGATIVE_REGULATION_OF_TRANSCRIPTION_FROM_RNA_POLYMERASE_II_PROMOTER_IN_RESPONSE_TO_STRESS
PEPTIDYL_SERINE_AUTOPHOSPHORYLATION
NEGATIVE_REGULATION_OF_INTERLEUKIN_6_PRODUCTION
REGULATION_OF_TOLERANCE_INDUCTION
NAD_BIOSYNTHESIS_VIA_NICOTINAMIDE_RIBOSIDE_SALVAGE_PATHWAY
MICROGLIAL_CELL_PROLIFERATION
NEGATIVE_REGULATION_OF_IMMUNOGLOBULIN_PRODUCTION
REGULATION_OF_LEUKOCYTE_TETHERING_OR_ROLLING
POSITIVE_REGULATION_OF_STEROID_HORMONE_SECRETION
C21_STEROID_HORMONE_METABOLIC_PROCESS
NEGATIVE_REGULATION_OF_MAST_CELL_ACTIVATION
NEGATIVE_REGULATION_OF_MAST_CELL_ACTIVATION_INVOLVED_IN_IMMUNE_RESPONSE
SEGMENT_SPECIFICATION
RESPONSE_TO_INTERLEUKIN_6
PEROXISOME_PROLIFERATOR_ACTIVATED_RECEPTOR_SIGNALING_PATHWAY
REGULATION_OF_RETINOIC_ACID_RECEPTOR_SIGNALING_PATHWAY
NEGATIVE_REGULATION_OF_RETINOIC_ACID_RECEPTOR_SIGNALING_PATHWAY
NEGATIVE_REGULATION_OF_MAST_CELL_DEGRANULATION
REGULATION_OF_FAT_CELL_PROLIFERATION
NEGATIVE_REGULATION_OF_LIPID_BIOSYNTHETIC_PROCESS
POSITIVE_REGULATION_OF_SIGNAL_TRANSDUCTION_BY_P53_CLASS_MEDIATOR
TYPE_I_PNEUMOCYTE_DIFFERENTIATION
CYCLOOXYGENASE_PATHWAY
REGULATION_OF_LIPID_CATABOLIC_PROCESS
SEX_DIFFERENTIATION
MESODERMAL_CELL_FATE_SPECIFICATION
POSITIVE_REGULATION_OF_HEART_GROWTH
PROXIMAL_DISTAL_PATTERN_FORMATION
NEGATIVE_REGULATION_OF_CANONICAL_WNT_SIGNALING_PATHWAY
POSITIVE_REGULATION_OF_FATTY_ACID_OXIDATION
POSITIVE_REGULATION_OF_LIPID_CATABOLIC_PROCESS
MAMMARY_GLAND_EPITHELIUM_DEVELOPMENT
MALE_GENITALIA_DEVELOPMENT
HGPS6
HGPS10
HGPS9
HGPS18
HGPS16
HGPS15
HGPS11
HGPS5
HGPS12
HGPS3
HGPS4
HGPS8
HGPS1
HGPS2
HGPS14
HGPS7
WT10
WT5
WT6
WT8
WT1
WT2
WT7
WT9
WT3
WT4
B
KRAS_SIGNALING_DN
ALLOGRAFT_REJECTION
ESTROGEN_RESPONSE_EARLY
APICAL_JUNCTION
MYOGENESIS
PI3K_AKT_MTOR_SIGNALING
HEDGEHOG_SIGNALING
UV_RESPONSE_UP
MTORC1_SIGNALING
GLYCOLYSIS
NOTCH_SIGNALING
WNT_BETA_CATENIN_SIGNALING
MITOTIC_SPINDLE
HALLMARK_HYPOXIA
APOPTOSIS
INFLAMMATORY_RESPONSE
KRAS_SIGNALING_UP
EPITHELIAL_MESENCHYMAL_TRANSITION
UNFOLDED_PROTEIN_RESPONSE
MYC_TARGETS_V2
G2M_CHECKPOINT
ANGIOGENESIS
ESTROGEN_RESPONSE_LATE
CHOLESTEROL_HOMEOSTASIS
IL2_STAT5_SIGNALING
P53_PATHWAY
MYC_TARGETS_V1
E2F_TARGETS
TNFA_SIGNALING_VIA_NFKB
OXIDATIVE_PHOSPHORYLATION
REACTIVE_OXYGEN_SPECIES_PATHWAY
ANDROGEN_RESPONSE
DNA_REPAIR
ADIPOGENESIS
COAGULATION
APICAL_SURFACE
TGF_BETA_SIGNALING
INTERFERON_GAMMA_RESPONSE
UV_RESPONSE_DN
PROTEIN_SECRETION
SPERMATOGENESIS
INTERFERON_ALPHA_RESPONSE
FATTY_ACID_METABOLISM
PEROXISOME
HEME_METABOLISM
BILE_ACID_METABOLISM
COMPLEMENT
XENOBIOTIC_METABOLISM
PANCREAS_BETA_CELLS
IL6_JAK_STAT3_SIGNALING
5
0
5
10
15
Count
WT (NES>0)
HGPS (NES>0)
WT (NES<0)
HGPS (NES<0)

**Figure S2. Gene ontology (GO) biological processes affected in HGPS patients. (A)** Heatmap of top 100 differentially enriched GOBP GSEA pathways. For each pathway, Normalized Enrichment Score (NES) values were compared between HGPS samples and control samples using a two-group t-test. Pathways were ranked by p-value, and NES values. Top 100 pathways ($p<0.005$, $FDR<0.16$) were visualized as a clustered heatmap using Euclidean distance and the complete linkage clustering method.) **(B)** Number of HGPS or WT individuals with NES >0 or NES < 0 for 50 GSEA hallmark pathways. GSEA NES values were obtained using average gene changes in HGPS compared to WT.

### Figure S3

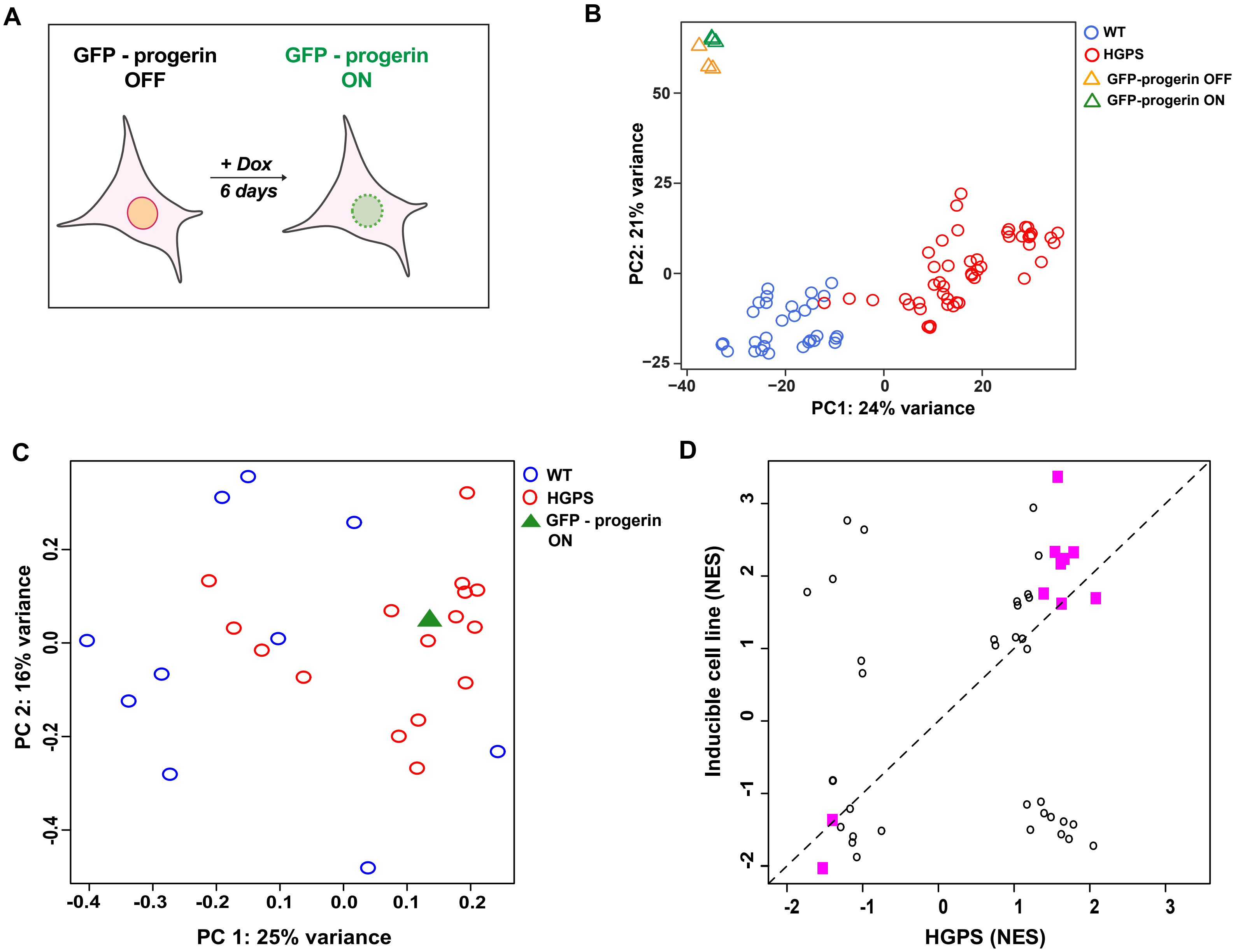

A
GFP - progerin OFF
GFP - progerin ON
+ Dox
6 days
B
WT
HGPS
GFP-progerin OFF
GFP-progerin ON
PC2: 21% variance
PC1: 24% variance
C
WT
HGPS
GFP - progerin ON
PC 2: 16% variance
PC 1: 25% variance
D
Inducible cell line (NES)
HGPS (NES)

**Figure S3. Comparison between primary patient fibroblasts and GFP-progerin inducible cell line.** **(A)** Schematic representation of stable human dermal fibroblasts cell line containing doxycycline-inducible GFP-tagged progerin. For analysis, GFP-progerin was expressed for 6 days. **(B)** PCA of the primary fibroblast and the GFP-progerin inducible cell line datasets used in this study. GFP-progerin cells show a largely different transcriptome compared to HGPS patients or control samples. The top 500 genes with the highest variability across samples were used for PCA plot. Orange triangles represent uninduced GFP-progerin cell line, green triangles represent GFP-progerin cell line where progerin was expressed for 6 days, blue dots represent wild type control samples, and red dots represent HGPS patients. All samples were analyzed in triplicate. **(C)** PCA plot using mean-centered NES values of 50 hallmark pathways in primary fibroblasts vs. GFP-progerin inducible cell line. Pathway scores of GFP-progerin cell line are more similar to HGPS than the WT samples. Blue circles represent healthy wild type control samples, red circles represent HGPS patients and green triangles represent GFP-progerin cell line where progerin was expressed for 6 days. **(D)** NES scatter plot for 50 GSEA pathways. HGPS NES were obtained using average HGPS gene changes compared to all WT samples for 50 hallmark pathways. Magenta squares indicate 10 pathways significantly affected ($padj < 0.05$) with consistent positive or negative trends in both GFP-progerin expressing cell line and in HGPS patients. Black circles indicate pathways affected only in HGPS or GFP-progerin expressing cells.

#### SUPPLEMENTAL TABLES

**Table S1.** HALLMARK PATHWAYS IN HGPS PATIENTS.

**Table S2.** GO_BP PATHWAYS IN HGPS PATIENTS. NES values for each patient are shown in each column.

**Table S3.** HALLMARK PATHWAYS IN GFP-PROGERIN CELLS.

**Table S4.** GO_BP PATHWAYS IN GFP-PROGERIN CELLS AND HGPS PATIENTS. NES values for each patient and GFP-progerin cell line are shown in each column.
